## Supplemental Material for "Reconfigurations of cortical manifold structure during reward-based motor learning"

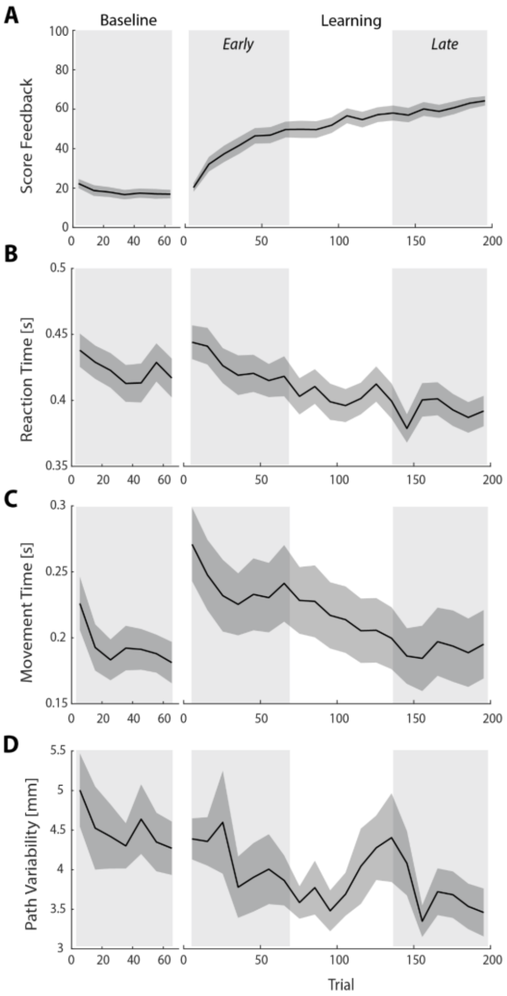


**Supplemental Figure 1: Behavioral measures of learning across the task.** (A-D) shows average participant reward scores (A), reaction times (B), movement times (C) and path variability (D) over the course of the task. In each plot, the black line denotes the mean across participants and the gray banding denotes +/- 1 SEM. The three equal-length task epochs for subsequent neural analyses are indicated by the gray shaded boxes.

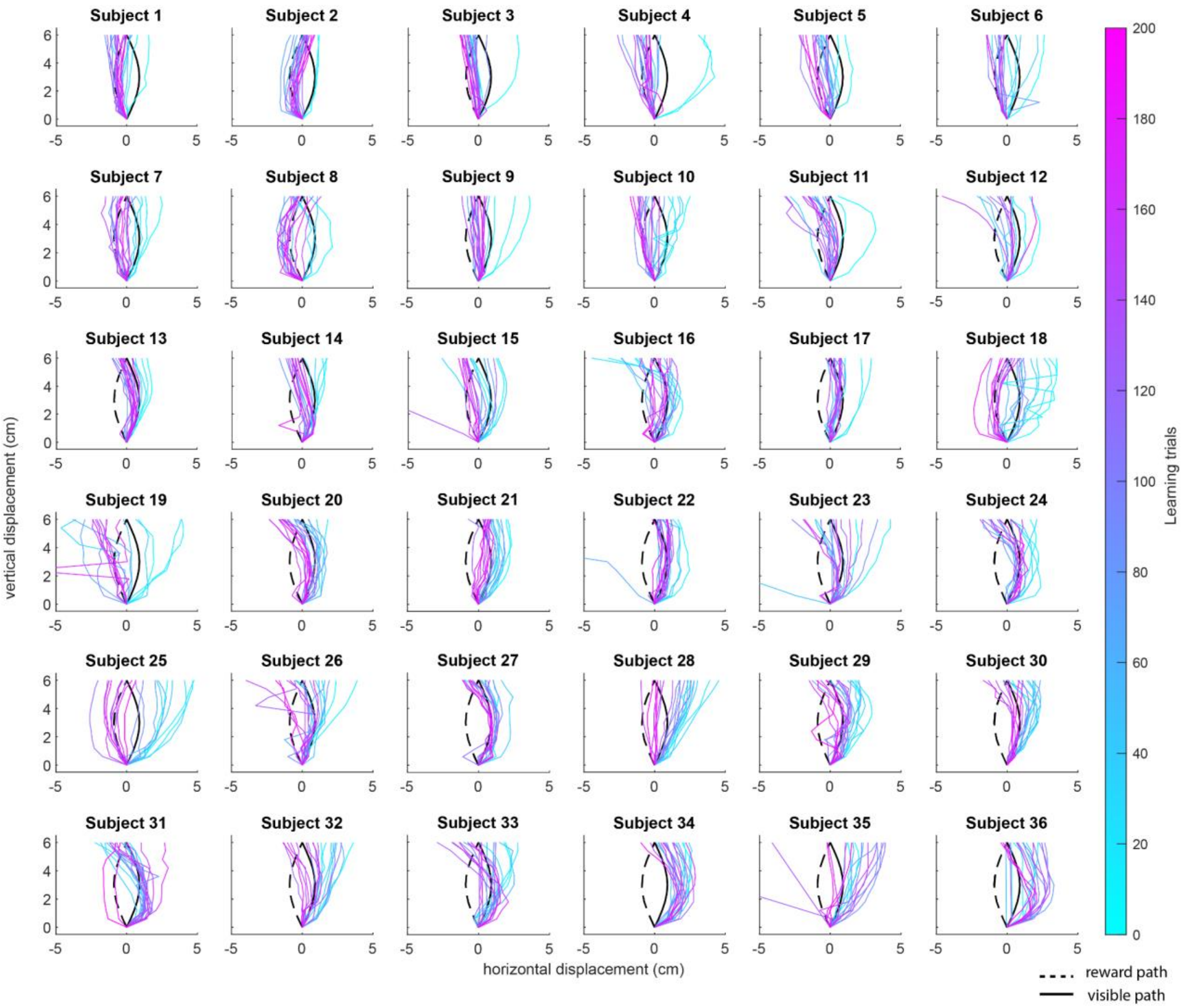


**Supplemental Figure 2: Variability in learning across subjects.** Plots show representative trajectory data from each subject (n=36) over the course of the 200 learning trials. Coloured traces show individual trials over time (each trace is separated by ten trials, e.g., trial 1, 10, 20, 30, etc.) to give a sense of the trajectory changes throughout the task (20 trials in total are shown for each subject).

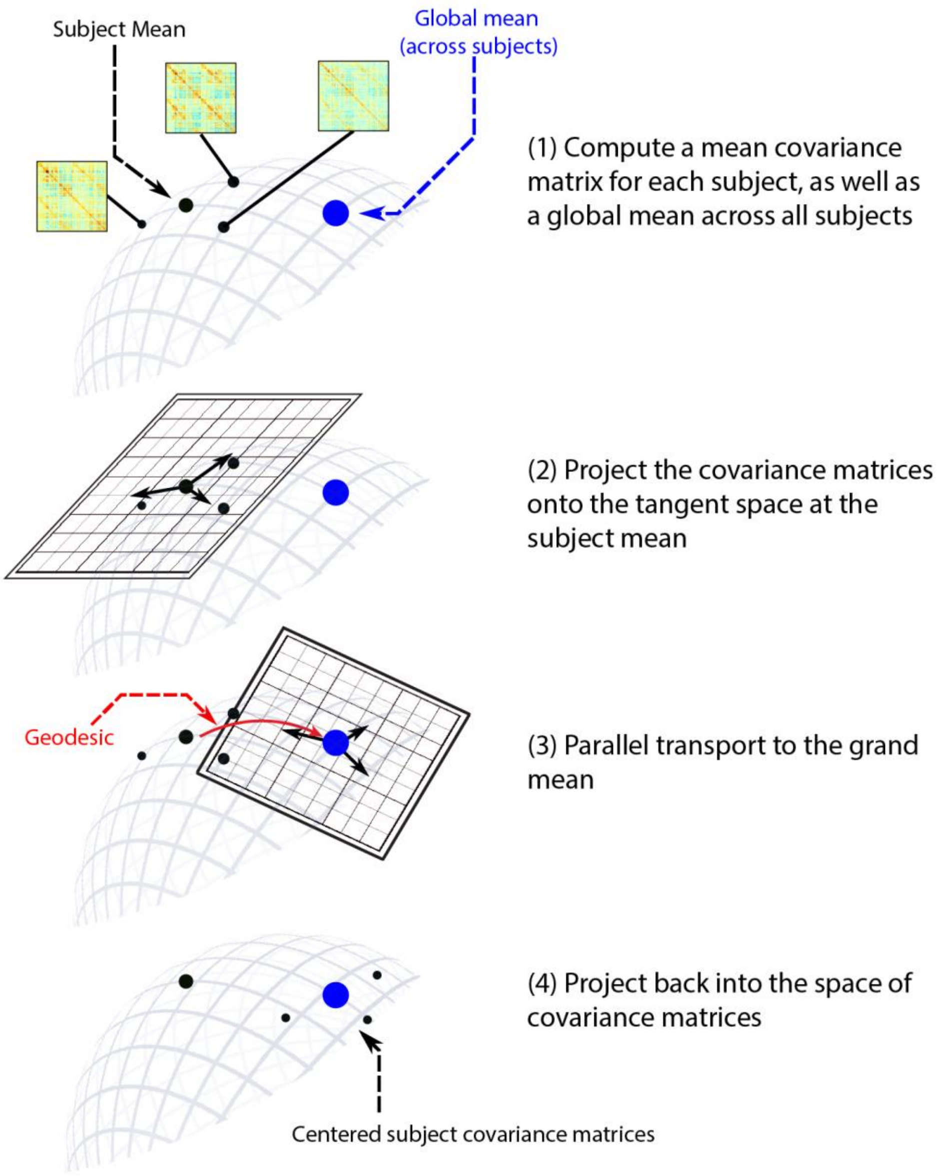


**Supplemental Figure 3: Overview of the Riemannian manifold centering approach.** To minimize the impact of significant individual variations in functional connectivity that might mask task-related changes, we used a Riemannian manifold approach to center all connectivity matrices. See steps #1-4 above for an overview. In short, each participant's covariance matrices were adjusted to share a common mean (equal to the overall mean covariance) to eliminate static individual differences that could potentially conceal task-related differences in functional connectivity. Since the space of covariance matrices is non-Euclidean, this adjustment cannot be achieved by simple subtraction. Instead, it requires calculating the difference between each covariance matrix and the corresponding participant mean (formally, a tangent vector) and then transporting this tangent vector to the overall grand mean to obtain a new covariance matrix that deviates from the grand mean in a comparable manner. For detailed computations involved in this process, please refer to the Supplementary Materials and Methods.

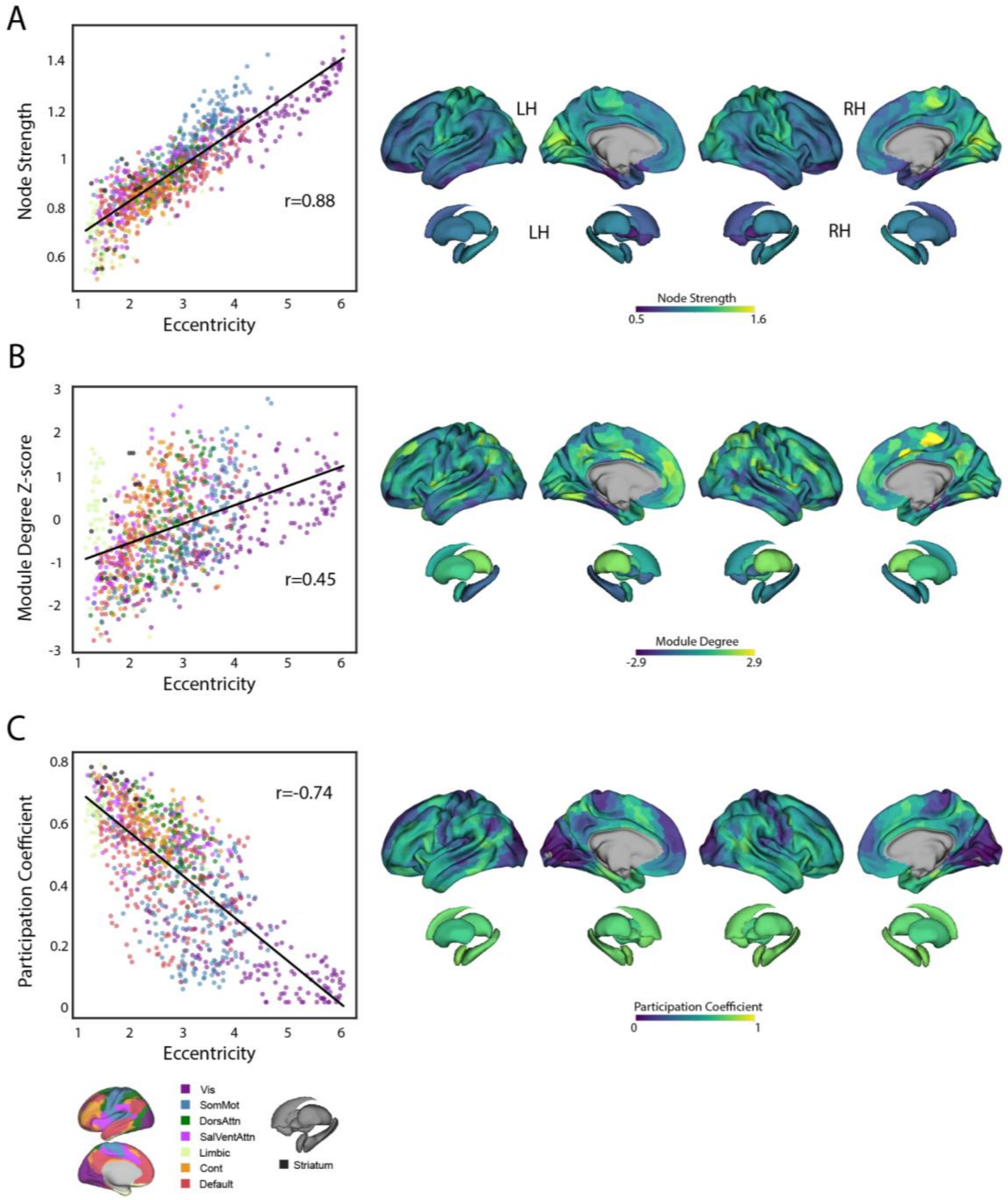


**Supplemental Figure 4: Functional connectivity properties that underlie manifold eccentricity.** (A-C) Different network properties of individual brain areas (derived from functional connectivity) and their correspondence to regional eccentricity. Left, scatterplots show the relationship between each functional connectivity measure and manifold eccentricity, with the line depicting a regression line of best fit to the cortical and striatal data. Right, brain plots show the maps of node strength, within-module degree z-score and participation coefficient, derived from the group-average Baseline connectivity matrix (i.e., reference connectivity matrix).

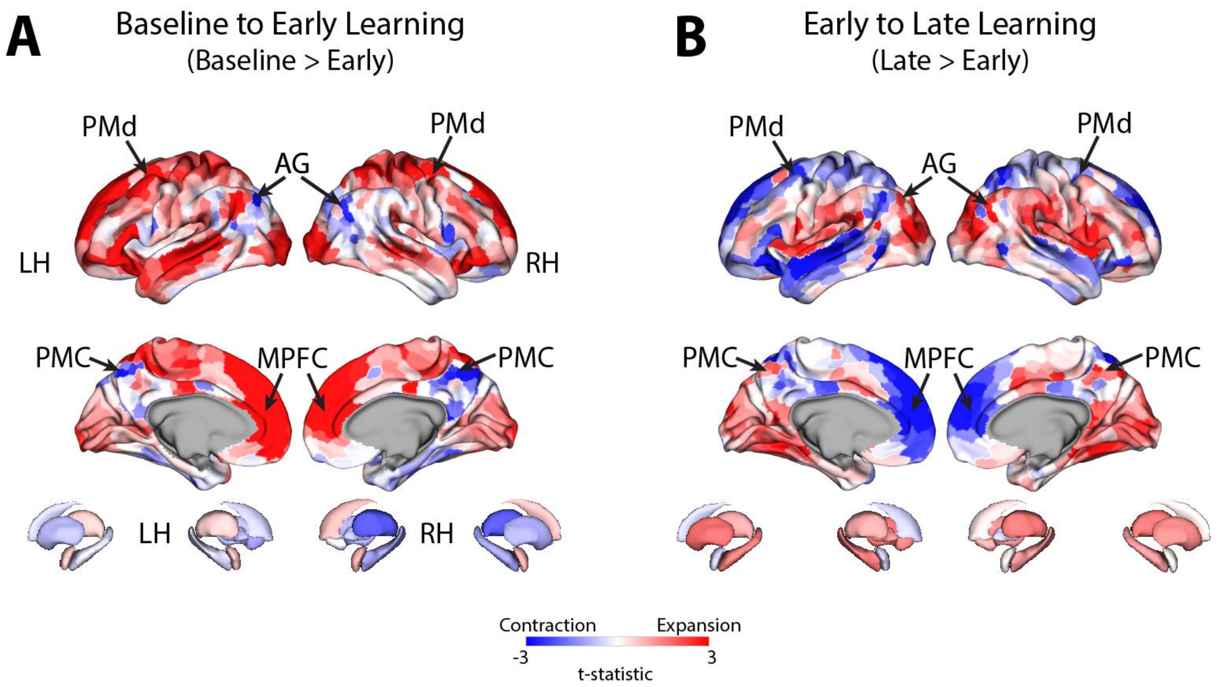


**Supplemental Figure 5: Unthresholded maps of changes in manifold structure during early and late learning.** (A & B) Pairwise contrasts of eccentricity between task epochs. Positive (red) and negative (blue) values denote increases and decreases in eccentricity (i.e., expansion and contraction along the manifold), respectively. The data shown in this figure is an unthresholded version of the data shown in main manuscript Figure 4.

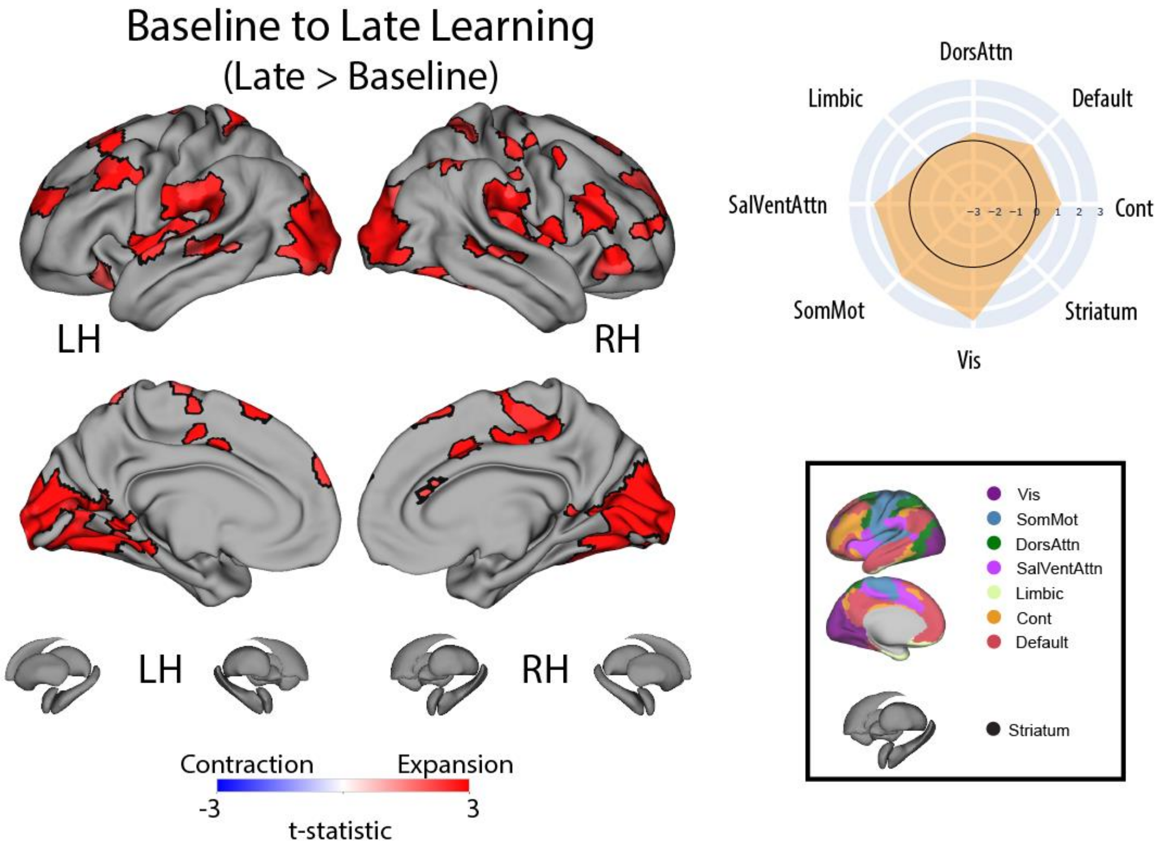


**Supplemental Figure 6:** Changes in manifold structure from Baseline to Late learning. Positive (red) and negative (blue) values show significant increases and decreases in eccentricity (i.e., expansion and contraction along the manifold), respectively, following FDR correction for region-wise paired t-tests (at $q<0.05$). The spider plot, at right, summarizes these patterns of changes in connectivity at the network-level (according to the Yeo networks). Note that the black circle in the spider plot denotes $t=0$ (i.e., no change in eccentricity between the epochs being compared). Radial axis values indicate t-values for the associated contrast.

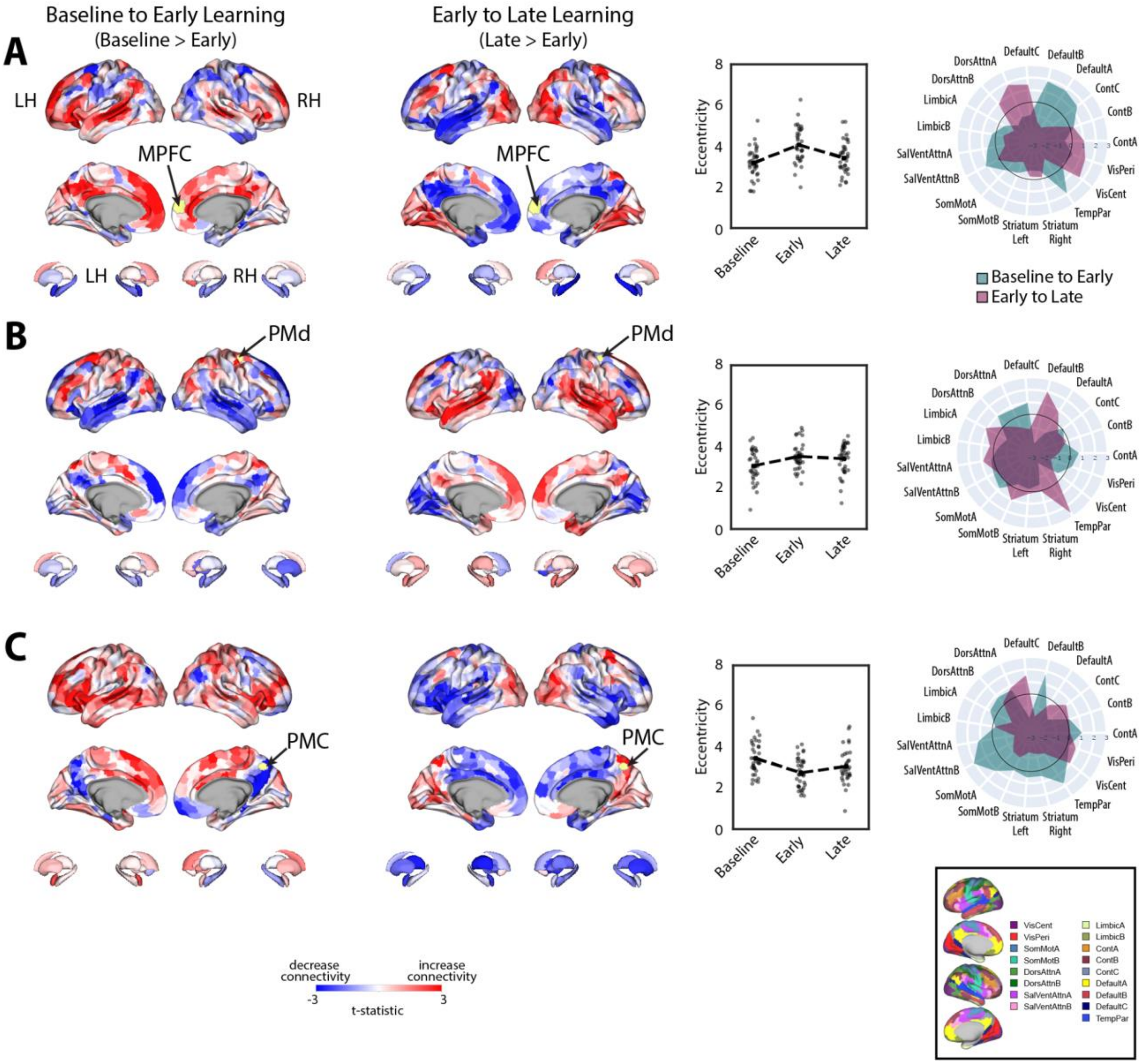


**Supplemental Figure 7: Patterns of connectivity changes that underlie manifold expansions and contractions for right-hemisphere regions.** (A-C) Connectivity changes for each seed region. Selected seed regions are shown in yellow and are also indicated by arrows. Positive (red) and negative (blue) values show increases and decreases in connectivity, respectively, from Baseline to Early learning (leftmost panel) and Early to Late learning (second from leftmost panel). Second from the rightmost panel shows the eccentricity of each region for each participant, with the black dashed line plot overlay showing the group mean across task epochs. Rightmost panel contains spider plots, which summarize these patterns of changes in connectivity at the network-level (according to the Yeo 17-networks parcellation (Yeo et al., 2011)). Note that the black circle in the spider plot denotes t=0 (i.e., zero change in eccentricity between the epochs being compared). Radial axis values indicate t-values for associated contrast.

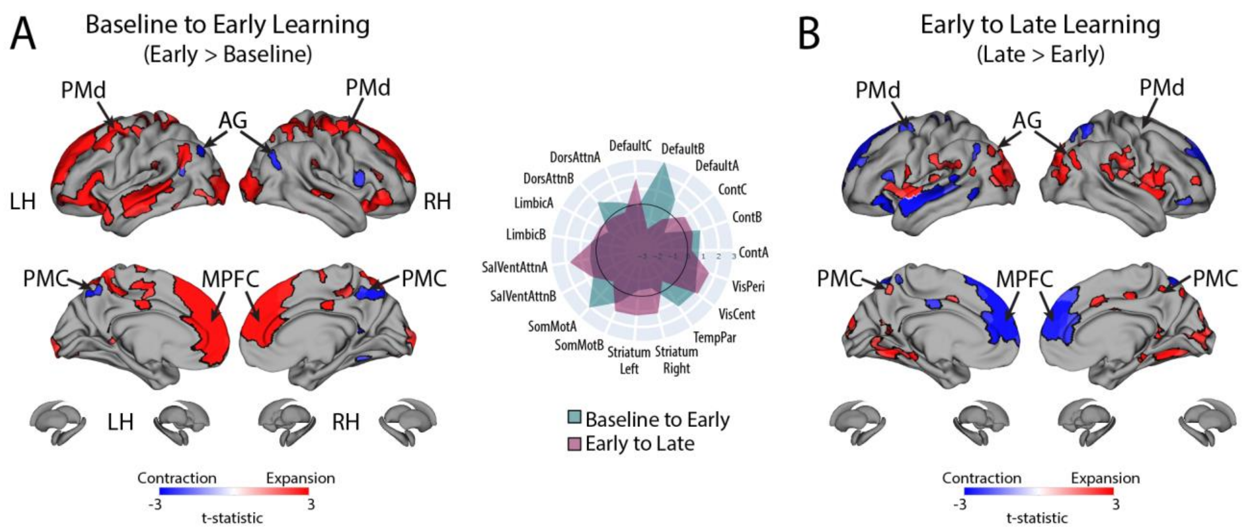


**Supplemental Figure 8: Changes in manifold structure during reward-based motor learning for the Yeo 17-network parcellation.** (A & B) Pairwise contrasts of eccentricity between task epochs. Positive (red) and negative (blue) values show significant increases and decreases in eccentricity (i.e., expansion and contraction along the manifold), respectively, following FDR correction for region-wise paired t-tests (at $q<0.05$). This data is the same as shown in Fig. 4 in the main manuscript. However, the spider plot, at center, summarizes these patterns of changes in connectivity at the 17-network-level (according to the Yeo networks, (Yeo et al., 2011)). Note that the black circle in the spider plot denotes $t=0$ (i.e., no change in eccentricity between the epochs being compared). Radial axis values indicate t-values for the associated contrast (see color legend).

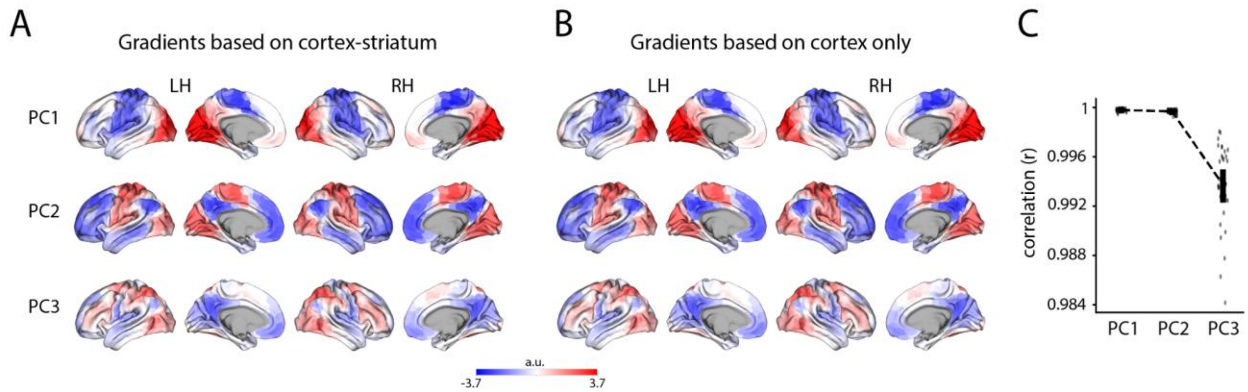


**Supplemental Figure 9: Derivation of cortical gradients did not depend on the inclusion of the striatum in the analysis.** (A) PCs 1-3 based on PCA decomposition of the group-average template baseline functional connectivity matrix, which included both cortical and striatal regions (as in Fig. 3A). (B) Same as in A, but based on a group-average template baseline functional connectivity matrix that did not include the striatal regions. (C) Whole-brain Pearson correlations between the data from A and B, for each PC. Vertical lines denote the mean +/-1 standard error. Single data points denote single subjects.
